# Selection bias in mutation accumulation

**DOI:** 10.1101/2021.08.03.454915

**Authors:** Lindi M. Wahl, Deepa Agashe

**Author notes:** **Author Contributions** LMW designed and conducted the study, and wrote the manuscript. DA contributed to the design of the analysis and wrote the manuscript. **Data Accessibility:** All of the simulation code used in this study is freely available at https://github.com/lmwahl/Selection_in_MA.

## Abstract

Mutation accumulation (MA) experiments, in which *de novo* mutations are sampled and subsequently characterized, are an essential tool in understanding the processes underlying evolution. In microbial populations, MA protocols typically involve a period of population growth between severe bottlenecks, such that a single individual can form a visible colony. While it has long been appreciated that the action of positive selection during this growth phase cannot be eliminated, it is typically assumed to be negligible. Here, we quantify the effect of both positive and negative selection in MA studies, demonstrating that selective effects can substantially bias the distribution of fitness effects (DFE) and mutation rates estimated from typical MA protocols in microbes. We then present a simple correction for this bias which applies to both beneficial and deleterious mutations, and can be used to correct the observed DFE in multiple environments. Finally, we use simulated MA experiments to illustrate the extent to which the MA-inferred DFE differs from the underlying true DFE, and demonstrate that the proposed correction accurately reconstructs the true DFE over a wide range of scenarios. These results highlight that positive selection during microbial MA experiments is in fact not negligible, but can be corrected to gain a more accurate understanding of fundamental evolutionary parameters.

## 1 Introduction

To estimate the rates, magnitude and spectra of spontaneous mutations – the raw materials for evolution (Dobzhansky, 1938) – mutations must be observed in the absence of selection. Experimental evolution under mutation accumulation (MA) thus serves as an essential approach for understanding evolutionary processes. MA experiments impose serial population bottlenecks, ideally with a single randomly-selected individual allowed to reproduce in each generation (Mukai, 1964). The resulting decrease in effective population size minimizes the impact of selection and the lineage accumulates all but lethal mutations, whose impacts on fitness can be examined in comparison with lineages evolving under selection. MA experiments have thus become an essential tool for evolutionary genetics (Halligan and Keightley, 2009, Katju and Bergthorsson, 2018). Such experiments are used to estimate key parameters in population genetics such as the distribution of fitness effects (DFE), mutation rates, and mutation spectra, and to shed light on the molecular mechanisms underlying changes in these parameters (Halligan and Keightley, 2009, Katju and Bergthorsson, 2018). In turn, these estimates underlie our understanding of a wide range of evolutionary phenomena including ageing, the evolution of mutation rates, the likelihood of adaptation, and the rate of mutational meltdown.

Enforcing population bottlenecks of a single reproductive individual (or reproductive pair) in each generation is not difficult to implement for large organisms such as worms and flies. For microscopic species such as bacteria and single-celled yeast, however, bottlenecks in which each colony on an agar plate is initiated by a single cell are possible, but cannot be imposed in every generation. This is because acquiring visible colonies on agar plates necessitates multiple generations of growth between transfers (i.e. between bottlenecks). The number of generations typically varies around 15-30 rounds of cell division for yeast or bacterial colonies (Foster et al., 2015, Joseph and Hall, 2004, Zeyl and DeVisser, 2001), allowing beneficial mutations to spread within the colony (Gralka et al., 2016).

Selection during growth between bottlenecks can thus introduce biases in MA results, even with a bottleneck of one individual at each transfer. As illustrated in Figure 1A, if we compare a beneficial and deleterious mutation that occur at the same time during colony growth, the beneficial mutation will, on average, be over-represented in the colonies that are formed on the subsequent agar plate. Such biases can lead, for example, to an underestimation of the deleterious mutation rate (Gordo and Dionisio, 2005, Kibota and Lynch, 1996, Trindade et al., 2010). A recent simulation study also demonstrates that selection can substantially favour the observation of beneficial mutations during MA (Mahilkar et al., 2021). Although many studies acknowledge the potential selection bias during MA, it is relatively rarely quantified and corrected. Kibota and Lynch corrected for the effect of selection against deleterious mutations (negative selection) when estimating the deleterious mutation rate in *Escherichia coli* (Kibota and Lynch, 1996). The same correction method was also used in some subsequent studies (e.g. Zeyl and DeVisser (2001)). Trindade et al. (2010) likewise corrected for the effect of negative selection, but were able to relax the assumption that the distribution of deleterious mutations in the population would reach equilibrium between bottlenecks (Gordo and Dionisio, 2005). For the yeast *Saccharomyces cerevesiae*, Joseph and Hall (2004) used a bias correction proposed for the conceptually similar accumulation of somatic mutations during a multicellular organism’s lifespan (Otto and Orive, 1995). As a result, the maximum likelihood estimate of the fraction of beneficial mutations was reduced from as high as 38% to 5.75% after this correction. In general, these methods adjust estimates of underlying mutation rates, or the mean deleterious effect of mutations, for negative selection; the effects of positive selection have typically been assumed negligible or explored in simulations (Mahilkar et al., 2021, Trindade et al., 2010). None of these approaches allows a correction to the DFE itself.

**Figure 1:**
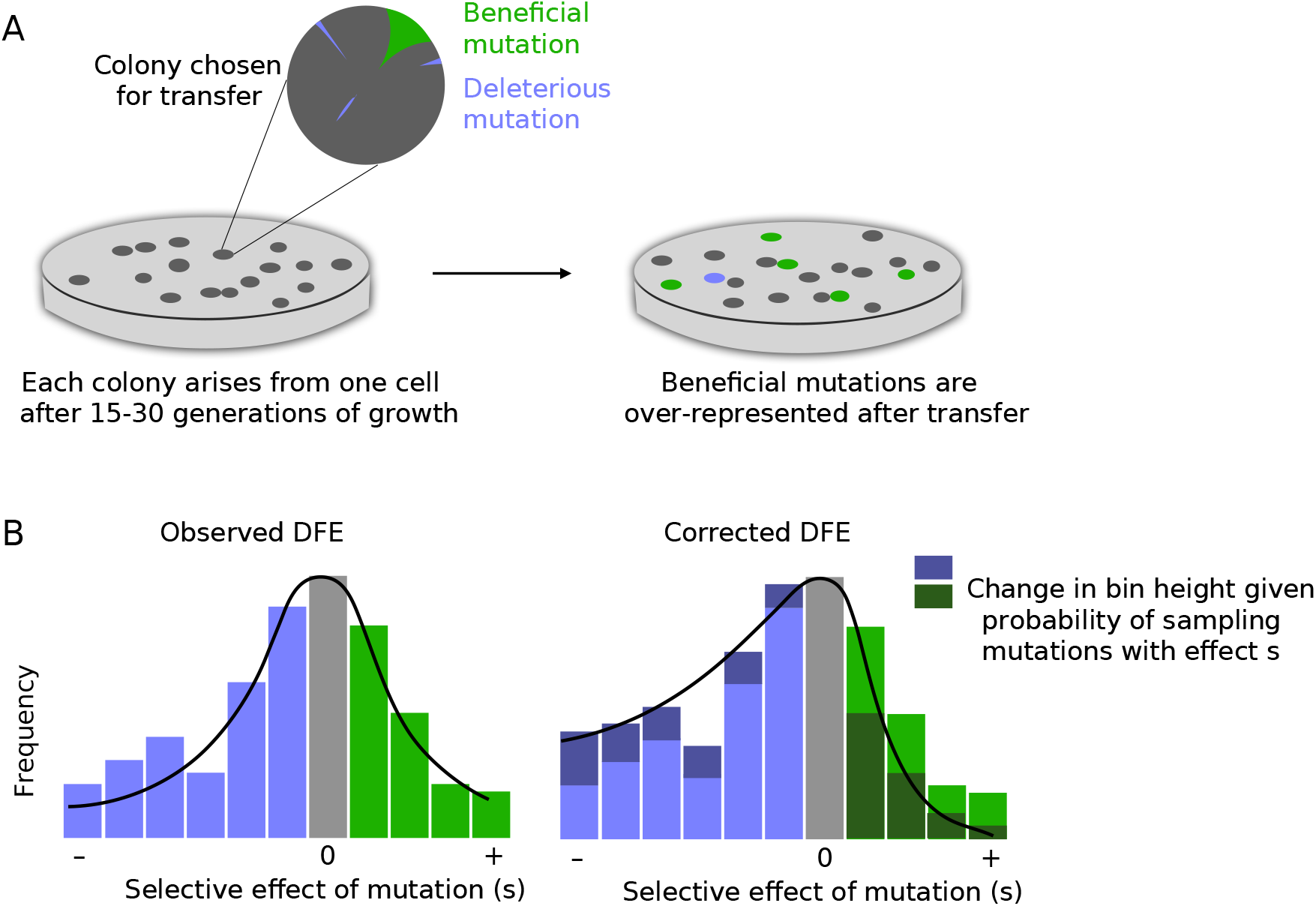
(A) Illustration of selection bias operating during colony growth between singlecell bottlenecks in microbial MA experiments. (B) Schematic showing the logic of our proposed correction for the distribution of fitness effects (DFE), to account for selection bias that would increase the observed frequency of beneficial mutations and reduce the frequency of deleterious mutations. The proposed correction increases or decreases the height of each fitness bin in the histogram by an amount that depends on its selective effect.

Here, we quantify the effects of both positive and negative selection during MA, showing why and to what extent DFEs are positively biased by the period of growth between serial bottlenecks. We provide a simple formula to correct observed DFEs for these effects. We also demonstrate that as long as the MA protocol isolates single mutations, selection has the same impact irrespective of the underlying mutation rate. Finally, we provide a method to correct for selection bias when the MA experiment is conducted in one environment and the DFE is measured in another environment. Our results thus offer a concrete and widely applicable way to derive more accurate inferences from experimental MA data.

## 2 Methods

We use a simple deterministic model to estimate the effects of selection in a typical mutation accumulation protocol. The protocol consists of repeated growth phases, during which a population with initial size *N*_0_ grows exponentially at rate *r* per unit time, such that the population size at any time is given by *N*(*t*) = *N*_0_*e*^*rt*^. For a pure birth process in bacteria, for example, we can express time in population doublings, such that *r* = ln 2 and the population size is *N* (*t*) = *N*_0_2^*t*^. In mutation accumulation protocols, experiments are typically designed to maintain *N*_0_ = 1. After a growth period of length *τ*, a sample of size *N*_0_ is taken from the population to found the next growth phase, and the process repeats.

We assume that mutations occur at a fixed rate *μ* per individual per unit time. This mutation rate will cancel out of the bias expression and in fact the results to follow are equivalent whether we assume mutations happen with a fixed probability per individual per unit time, or per replication. New mutations have selection coefficient *s*, which may be positive or negative, such that the Malthusian growth rate of a mutant lineage is *r*(1 + *s*).

In a Wright-Fisher population of a constant size, each new generation is formed by random sampling of a much larger number of offspring, and therefore new lineages are frequently lost to genetic drift. We assume that genetic drift has been minimized during the growth phase of the mutation accumulation protocol, that is, the chance that a new mutation fails to establish is negligible. This assumption could easily be relaxed for populations that do not experience a pure birth process during the exponential growth phase, by adding a survival term that depends on *s*.

At time *t* during the growth phase, new mutations occur at rate *μN*_0_*e*^*rt*^. Assuming no mutations are lost to drift, each mutation creates a lineage that grows to final size *e*^*r*(1+*s*)(*τ* – *t*)^ at time *τ*, when the population is sampled (i.e. a grown bacterial colony is transferred to a new agar plate).

To quantify the relative bias imposed by selection during the growth phase, we compare 110 the probability that a mutation with effect *s* is chosen during sampling to the probability that the same mutation would have been chosen if it had an effect *s* = 0. We do this simply by comparing lineage sizes at time *τ*, conditioned on the time *t* at which the mutation first occurred:

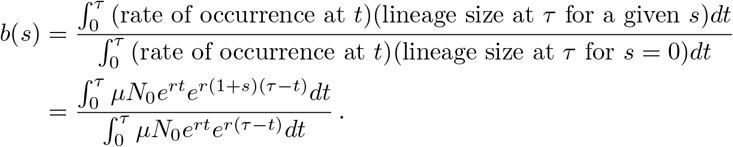

This is similar to the approach suggested by Kibota and Lynch (1996), who considered only deleterious mutations, whereas we treat both beneficial and deleterious mutations. We also note that Kibota and Lynch reduced the ancestor growth rate by the (deleterious) mutation rate, obtaining *N* (*t*) = *N*_0_*e*^(*r*–*μ*)*t*^. Here, we assume that the mutation rate is negligible compared to the growth rate. As a result of this simplifying assumption, our approach yields a compact expression for relative selection bias:

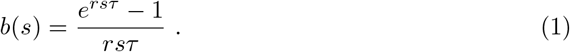

The relative selection bias, *b*(*s*), quantifies the factor by which a mutation with effect *s* is more or less likely to be chosen *during the population sampling after it first occurs*. If *N*_0_ > 1, this bias could accrue in further growth phases, and the overall chance of observing *s* would depend even more strongly on *s*. However when *N*_0_ = 1, the mutation either survives the first sampling after it occurs or it is lost; there is no chance to accrue a further benefit, for example, during a subsequent growth phase. Thus, for mutation accumulation protocols in which a single individual is chosen to be propagated, the overall bias due to selection is simply given by *b*(*s*).

The length of the growth phase, *τ*, in Equation 1 can be provided in any convenient time 129 units, as long as the Malthusian growth rate of the ancestor and mutant are given by *r* 130 and *r*(1 + *s*) per unit time respectively. Again, if we measure time in units of population 131 doublings, *r* = ln 2 and the selection bias for (or against) a mutation with effect *s* is given 132 by:

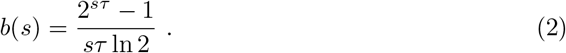

Equations 1 and 2 suggest several conclusions which we will confirm in simulation studies described below. It is clear that *b*(0) = 1, that is, neutral mutations have no increased or decreased chance of being selected. However *b*(*s*) > 1 for beneficial mutations, and *b*(*s*) < 1 for deleterious mutations, as expected. We can also see directly from this expression that bias will increase with *τ*, the number of generations during each growth phase. Finally, as mentioned previously, the mutation rate drops out of the expression for *b*(*s*); the selection bias is independent of the mutation rate.

How would we use these results to correct an observed distribution of selective effects of mutations (DFE)? An important point to note here is that this bias does not affect the fitness of the observed mutations (the *s* values remain unchanged), but affects the probability that they are observed during the MA experiment. Thus, the correction is applied to the DFE, not to individual fitness observations. The most straightforward way to to do this is to correct binned fitness data (a histogram), which are commonly used to visualize DFEs.

In particular, suppose we have a DFE described by *n* bins, when bin *i* is centred at selective effect *s*_*i*_, *i* = 1*..n*. To correct for selection bias, we re-weight the histogram such that events that we were more likely to observe because of selection are given less weight (and vice versa), as illustrated in Figure 1B. If *b*(*s*_*i*_) is the bias associated with the bin centre *s*_*i*_, we simply re-weight that bin by *w*_*i*_ = 1/*b*(*s*_*i*_). Letting *f*_*i*_ denote the observed frequency of mutations in bin *i*, the corrected frequency 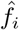 is given by re-weighting, and then re-normalizing so that the corrected frequencies sum to one:

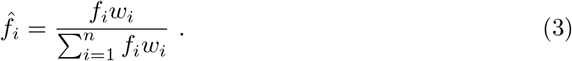

A useful way to think about this correction is that usually, we build a histogram by stacking similar fitness measures in vertical bins, in which we give a height of one to each fitness measurement in the bin (and then, optionally, normalize so that the bins sum to one). Here, we create a histogram in the same way, but in order to correct for selection, we give a height of *w*_*i*_ to each of the measurements in bin *i*.

### 2.1 Correcting the DFE in new environments

When fitness is also measured in a different environment than the environment in which mutations were first observed, correcting for selection bias is more subtle. For example, in Sane *et al.* (2018), 80 *E. coli* strains that each differed by a single distinct mutation from their ancestor were identified in an MA protocol in one environment, and the fitness of these 80 strains was then measured both in the original environment (“MA environment”) and in eleven other environments with different carbon sources (“new environments”).

In this case, we would like to assign a weight to each strain, based on the bias in observing that mutation. Intuitively, it makes sense that if fitness measurements in the new environment are perfectly correlated with fitness in the MA environment, we should apply the same correction, Equation 3, above. However if fitness in the new environment is completely *uncorrelated* with fitness in the MA environment, the set of mutations collected in the MA environment should provide a random sample of fitness values in the new environment, and we should therefore apply no correction to the DFE measured in the new environment. We can use the correlation coefficient as a simple heuristic to move between these two extremes.

In particular, suppose strain *m* has selective effect *s*_*m*_, measured in the MA environment. We compute *ω*_*m*_ = 1/*b*(*s*_*m*_), where *ω*_*m*_ represents the weight that strain *m* should be given in the DFE.

Suppose the fitness of these strains is then measured in a new environment. Let *ρ* be the correlation coefficient between the two sets of fitness measurements. To build a DFE for the new environment, corrected for selection, we build a histogram of selective effects, measured in the new environment. But instead of giving each observation a height of unity in the histogram, we give each observation a height of 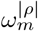. When │*ρ*│ = 1 this procedure will be equivalent to Equation 3, and the same correction will be applied in both the MA and the new environment. In contrast when *ρ* = 0, this procedure will give every measurement a weight of one; no correction is applied in the new environment. For intermediate correlation coefficients, the degree to which the correction is applied scales smoothly between these two extremes. The absolute value of the correlation coefficient is necessary because we always want to reduce the height for mutations that were over-represented during the MA, whether those tend to be on the left or the right of the DFE in the new environment.

Mathematically, suppose strain *m* has selective effect *s*_*m*_ in the MA environment and effect 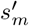 in the new environment. If we use bin^′^(*m*) to denote the histogram bin to which the fitness in the new environment, 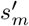, belongs, the corrected frequency of bin *i* in the new environment is given by:

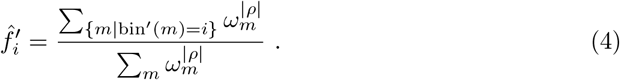

As noted above, this is a relatively simple approach that scales the degree of correction smoothly between no correction (at *ρ* = 0) and full correction (at *ρ*=1,−1), but other approaches (such as scaling the bin heights linearly between 1 and *ω*_*m*_) would also be reasonable.

### 2.2 Simulations

We performed stochastic simulations of a mutation accumulation (MA) protocol with

*N*_0_ = 1 to confirm our analytical predictions. In brief, each simulation begins with a single individual with absolute fitness *W*_0_ = 2, that is, an individual that contributes, on average, two individuals to the next generation. We model *τ* population doublings, and then choose one individual at random from the population to found the next colony (a ‘transfer’). We repeat this process in a large number of *in silico* lines, in parallel, and for a large number of transfers.

At each birth event, a mutation occurs with probability *μ*, and if so, the selective effect of the mutation, *s* is drawn randomly from a probability distribution; *s* can be positive or negative. The new individual’s fitness is then given by the parental fitness multiplied by (1 + *s*).

An individual with fitness *W*_*i*_ should contribute, on average, *W*_*i*_ individuals to the next generation. A poisson-distributed random number with mean *W*_*i*_ ≈ 2, however, has substantial weight at zero, and thus describes a population with a substantial death rate; this is not an appropriate model for most experimental MA protocols. To model a pure birth process in which new lineages are not lost, each individual in our simulation contributes one individual to the next generation (i.e. the parent bacterium continues to the next generation), plus a poisson-distributed number of offspring with mean *W*_*i*_ – 1.

To model the distribution of selective effects, in many of the examples shown below we used a shifted, reflected gamma distribution (Bataillon and Bailey, 2014, Eyre-Walker and Keightley, 2007, Martin and Lenormand, 2006, Nielsen and Yang, 2003). In particular, if Г_(*k,θ*)_(*s*) denotes a gamma distribution with shape parameter *k* and scale parameter *θ* (thus mean *kθ*), the probability density for selective effects is given by:

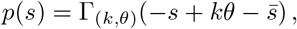

for 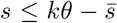, while *p*(*s*) = 0 for values of 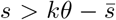. Thus the gamma distribution is reflected about the *y*-axis and then translated to have mean 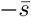; distributions with higher values of *k* are narrower.

To test the robustness of our results to the shape of the underlying DFE, we manipulated the parameters of the gamma distribution above, and also tested a different analytical form for the underlying DFE, given by a mixture of exponential distributions (Böndel et al., 2019, Mahilkar et al., 2021). For this second case, we take

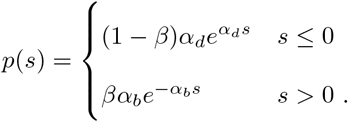

In this “double exponential” DFE, mutations with positive effect occur with probability *β*, and are distributed exponentially with mean 1/*α*_*b*_. Mutations with negative effect occur with probability 1 – *β* and are likewise distributed exponentially, but with mean −1/*α*_*d*_. This analytical form allows us to independently manipulate the beneficial fraction, as well as the beneficial and deleterious effect sizes.

On a practical note, the number of generations between transfers can be as large as *τ* = 30 in experimental protocols. The population size thus grows to hundreds of millions of individuals, necessitating an approach that does not track each individual and fitness value. Instead, the simulation code tracks the number of individuals in each fitness bin, where for example we divide the range of fitness values from *W* = 0 to *W* = 3 into 1500 bins.

We also simulated experimental scenarios in which the fitness of strains isolated during MA is measured in different environments, assuming that fitness in each environment is correlated to fitness in the MA environment according to a known correlation coefficient, *ρ* (see SI). We can thus manipulate the degree of correlation between fitness values measured in the two environments, apply the correction described in Equation 4, and compare corrected DFEs to a simulated “true” DFE in that environment.

Finally, the goal of an MA protocol, in principle, is to measure the fitness effects of *single* mutations. To achieve this in the simulations, for each fitness bin we not only track the total number of individuals in the bin, but we also count how many of those individuals differ by exactly one mutation from the founder of the current colony. When a random individual is chosen to found the next colony, it is thus possible to reconstruct whether this individual differs from the previous founder by zero, one or more than one mutation. As in our experimental protocol (Sane et al., 2018), in the simulated data only individuals that differ by a single mutation from their colony founder are included in the DFEs.

Unless otherwise noted, for each simulated MA experiment, 1000 independent lines were simulated for between 5 and 27 generations of growth between transfers. A single cell was chosen at each transfer to start the growth of the next colony. The number of transfers per experiment varied with mutation rate to obtain approximately 1000 single mutations in each simulated DFE. The default parameters for mutation were *μ* = 0.00043 mutations per genome per generation (the empirical estimate in Sane et al. (2018)), with *p*(*s*) defined using *k*=10, *θ*=0.05 and 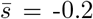. In generating summary statistics, we considered neutral mutations to be those within the range −.05 < *s* < 0.05, corresponding to the experimental error in fitness measures in Sane et al. (2018).

## 3 Results

Figure 2 shows the strength of the MA selection bias and how it affects the inferred DFE. Panels A and B illustrate the unexpectedly large magnitude of this effect; if the MA protocol includes a growth phase of 20-30 generations, for example, a large-effect beneficial mutation with *s* = 0.1 is *5 or 10 times more likely* to be observed than a neutral mutation that occurs with the same mutation rate. For any underlying DFE (black lines in panels C through F), selection substantially reduces the chance that a deleterious mutation is observed, consistent with established predictions and approaches (Hall and Joseph, 2010, Kibota and Lynch, 1996, Trindade et al., 2010), while also increasing the chance that a beneficial mutation is observed. For a gamma-distributed DFE, the overall effect (panel D) is a gradual change in the shape of the DFE, and a shift toward more positive values. All of these changes are exacerbated by longer periods of growth during the MA protocol.

**Figure 2:**
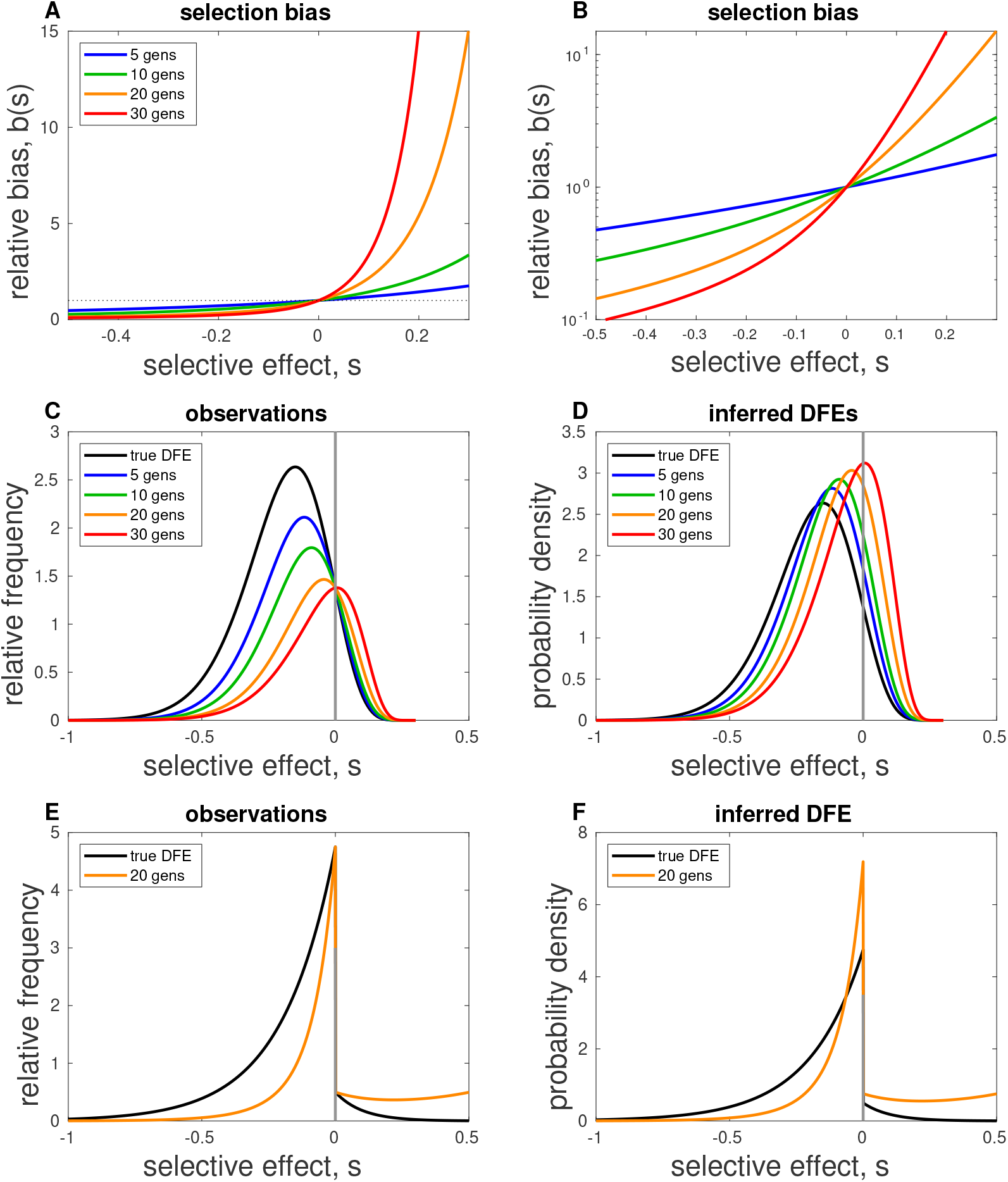
Effects of selection bias. (A) The magnitude of the relative bias, *b*(*s*) as a function of the selection coefficient, *s*; lines vary the number of generations of growth between transfers. To aid visualization when *s* < 0, panel (B) shows the same data on a semilog scale. (C) Predicted observations in a MA experiment, i.e. the product of the “true” DFE and the selection bias. Given an underlying DFE (black), selection reduces the chance that a deleterious mutation is observed, while increasing the chance that a beneficial mutation is observed. (D) Results from panel (B), normalized such that the area under each curve is unity. When the observed distribution is renormalized to infer a DFE, the net effect is a gradual change in shape and a beneficial shift in the inferred DFE. (E) As in panel (C) for 20 generations of growth, but in this case the underlying DFE was a mixture of two exponential dis2tr3ibutions. (F) Results from (E), normalized to show the inferred DFE. Parameter values: (C and D) *k*=10, *θ*=0.05, 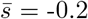; (E and F) *β* = .05, *α*_*d*_ = 5, *α*_*b*_ = 10.

In panels (E) and (F) of Figure 2, we illustrate the same effects using an underlying DFE composed of a mixture of exponential distributions. For visual clarity, we show results for *τ* = 20 generations of growth between transfers, a typical value in MA in yeast (Farlow et al., 2015, Lang et al., 2013, Zhu et al., 2014). For the gamma distribution, the fraction of mutations that are beneficial (*s* > 0) is 8% in the true DFE, and rises to 25% in the inferred DFE; for the exponential distribution, a beneficial fraction of 5% rises to 18% in the inferred DFE. We also note in panels (E) and (F) that selection can compensate or in fact over-compensate for the decay in the right tail of the DFE, resulting in an inferred DFE that is truncated on the right (see Rokyta et al. (2008) for a discussion of right-truncated DFEs obtained under strong selection).

In Figure 3, we show results of simulated MA experiments; the underlying DFE from which mutations were randomly drawn during the simulated MA was fixed (black lines in panels A, B and C), while the number of generations between transfers was varied. Observed DFEs were collected from the simulated data (purple histograms), and are increasingly different from the underlying DFE for longer growth phases. After correction using Equation 3, however, the DFEs (cyan) show excellent agreement with the true DFE. In the lower panels, we illustrate the fraction of the DFE that is beneficial, neutral or deleterious, and the mean selective effect of mutations in each of those three categories. These summary statistics can change dramatically with the length of the growth phase, but show excellent agreement with the true values from the underlying DFE once the correction has been applied to the histogram.

**Figure 3:**
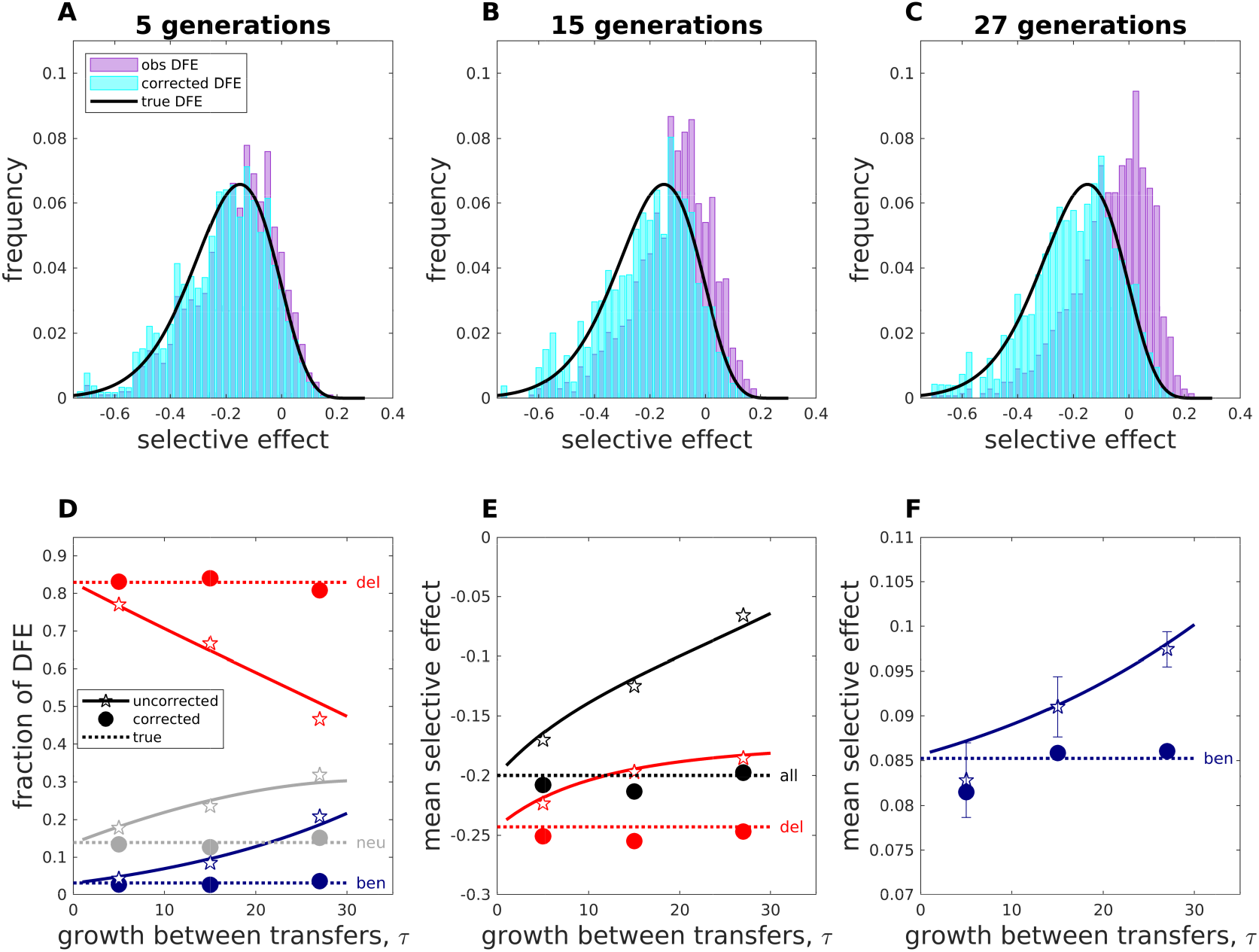
Selection bias and the length of the growth phase. Top row: Results of simulated MA experiments in which mutations are randomly drawn from the “true” DFE (black). Observed DFEs (purple) are shown for *τ* = 5 (panel A), 15 (panel B) or 27 generations of growth between transfers. Each observed DFE was corrected as described in Equation 3; corrected DFEs (cyan) show excellent agreement with the underlying DFE. Bottom row: The effect of selection on DFE summary statistics. If uncorrected, the fraction of the observed DFE that is deleterious (*s* < −0.05), neutral (−0.05 < *s* < 0.05), and beneficial (*s* > 0.05) varies with the number of generations of growth between transfers (solid lines, panel D). Likewise, the mean selective effect of all observed mutations, observed deleterious mutations and observed beneficial mutations all shift toward more positive values as *τ* increases (panels E and F). In the bottom row: stars show uncorrected simulation results, circles show corrected simulation results. Solid lines show theoretical expectations for uncorrected data; dotted lines show results given the “true” DFE from the top row. Error bars (± 1 standard deviation) were estimated from 1000 bootstrapped samples of each set of observed (simulated) *s* values. Error bars in panels D and E are similar to symbol heights and omitted for clarity.

Figure 4 shows similar results, but in this case the length of the growth phase is fixed at 20 generations, while the mutation rate in the simulated MA experiment is varied. For illustration, we used mutation rates estimated in MA experiments in *E. coli* (Sane et al., 2018, 2020), along with a higher mutation rate to study mutation rates that varied in total by factor of ≈ 20 in panels (A) through (C), and by a factor of 1000 in panels (D) through (F). While selection bias affects each of these DFEs due to the 20 generations of growth in the MA protocol, we find that the effects of selection on the DFE are independent of the mutation rate. Correction for selection bias once again accurately reproduces the underlying DFE and its associated summary statistics.

**Figure 4:**
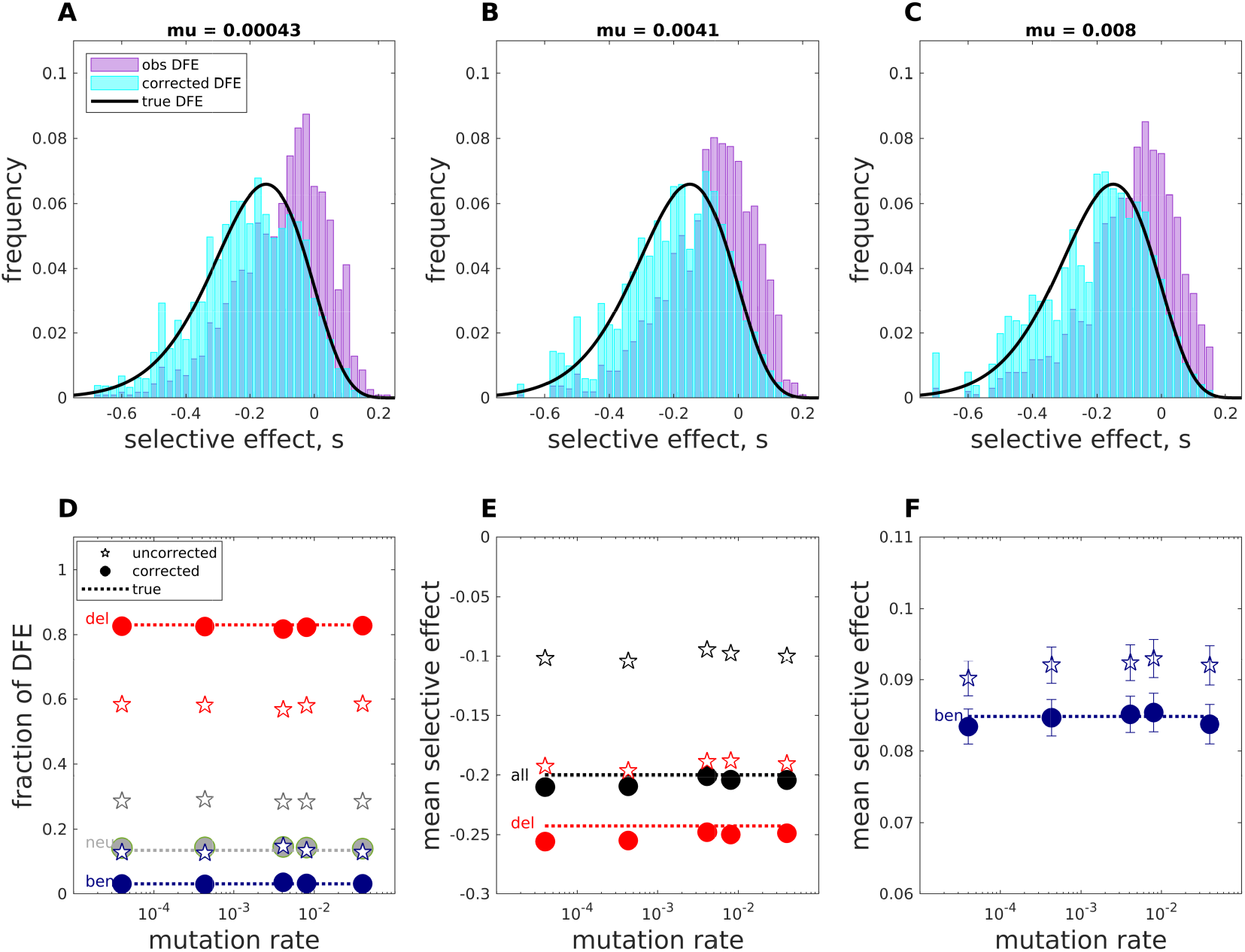
Selection bias and the mutation rate, *μ*. Top row: Results of simulated MA experiments in which mutations are randomly drawn from the “true” DFE (black). Observed DFEs (purple) are shown for *μ* = 0.00043 (panel A), 0.0041 (panel B), or 0.008 (panel C) mutations per genome per generation, with *τ* = 20 generations of growth between transfers in each case and sufficient independent lines to reach 1000-2000 individual mutations in each experiment. Each observed DFE was corrected for 20 generations of growth as described in Equation 3; corrected DFEs (cyan) show excellent agreement with the underlying DFE. Bottom row: DFE summary statistics are not sensitive to the mutation rate. (D) The deleterious (*s* < −0.05), neutral (−0.05 < *s* < 0.05), and beneficial (*s* > 0.05) fractions, for mutation rates varying across 3 orders of magnitude. Stars show uncorrected simulation data, circles show the corresponding simulation results after correction. Lines show theoretical expectations, given the true DFE shown in the top panels. (E) The mean selective effect of all observed mutations and the mean effect of observed deleterious mutations (*s* < −.05) are likewise insensitive to the mutation rate. (F) In the simulated data, the mean effect of observed beneficial mutations (*s* > 0.05) shows a tendency to shift toward more positive values at high mutation rates. Error bars (± 1 standard deviation) were estimated from 1000 bootstrapped samples of each set of observed (simulated) *s* values. Error bars in panels D and E are similar to symbol heights and omitted for clarity.

In Figure 5, we show results after varying the shape of the underlying DFE in the simulated MA experiment. Corrected DFEs show good agreement with the underlying DFE, even for relatively narrow or broad distributions, and across both analytical forms of the underlying DFE. We note that correcting for selection has a much larger effect in broader DFEs (panels A and C), due to the larger expected magnitude of *s*. In the lower panels, we show the fraction of the uncorrected DFEs that would be observed to be beneficial, neutral or deleterious, along with the corrected results; again, the correction has a much larger effect for broader DFE distributions. As expected, the observed beneficial fraction always increases, and the deleterious fraction decreases, with longer growth phases. Interestingly though, the neutral fraction can increase or decrease, depending on the shape of the underlying DFE (and the threshold used to define neutrality, here ± 5%).

**Figure 5:**
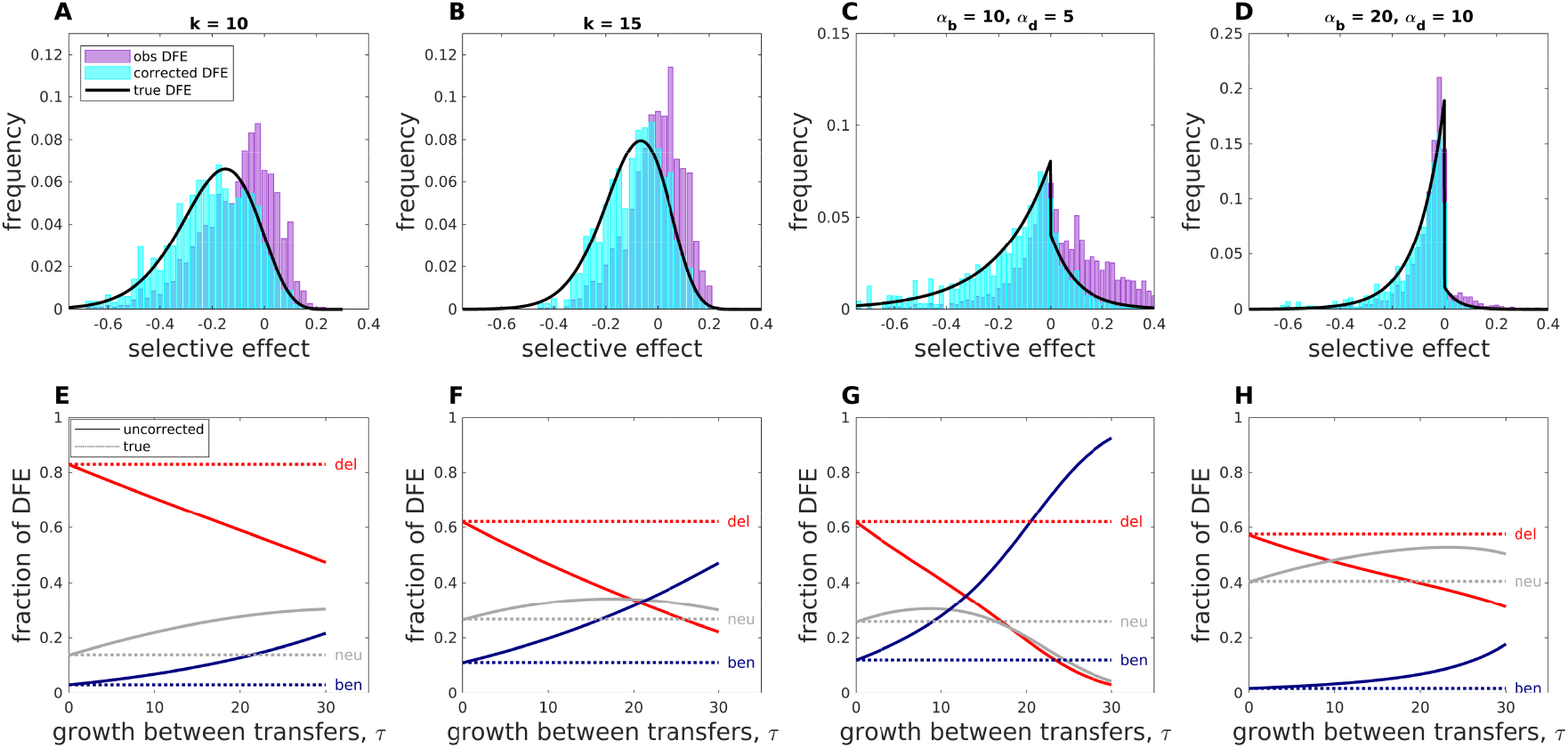
Selection bias and the shape of the DFE. Top row: Results of simulated MA experiments in which mutations are randomly drawn from the “true” DFE (black), which varies in each column. Observed DFEs (purple) are shown for the reflected gamma distribution with *k* = 10, 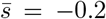 (panel A) or *k* = 15, 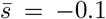 (panel B), with *θ* = 0.5/*k* in both cases. Analogous results using the double exponential distribution are shown with *β* = .2, *α*_*b*_ = 10, *α*_*d*_ = 5 (panel C) or with *β* = .05, *α*_*b*_ = 20, *α*_*d*_ = 10 (panel D); *τ* = 20 generations of growth between transfers in all four cases. Each observed DFE was corrected for 20 generations of growth as described in Equation 3; corrected DFEs (cyan) show excellent agreement with the underlying DFEs. (E through H): Solid lines plot DFE fractions that would be observed given the true DFE shown in the corresponding top panels; dotted lines show DFE fractions after correcting for selection. As *τ* increases, selection reduces the deleterious fraction and increases the beneficial fraction for all three underlying DFEs. Note however that for some DFE shapes, the neutral fraction can either increase or decrease with *τ*.

Finally, in Figure 6, we show an example of the application of these corrections to experimental data. In brief: 80 individual mutations were collected during an MA experiment with *E. coli* growing on a rich growth medium (Luria Bertani agar, LB); the fitness of this set of mutations was then measured in LB as well as different minimal media containing various carbon sources (Sane et al., 2018). The top panels of Figure 6 show the experimentally observed DFEs in LB and two other carbon sources (purple), along with the same DFEs after correction (cyan) using Equation 3 (panel A) or Equation 4 (panels B and C, note that the two histograms overlap completely in C). The lower panels show the effect of the correction on the summary statistics for these DFEs. We note that correction for selection bias reduces the beneficial fraction and increases the deleterious fraction of the DFE when measured in the MA environment. In particular, the experimental estimate of the beneficial fraction fell from 40% to 5.8% after correction, and thus the estimate of the rate of beneficial mutation fell from 1.7 × 10^−4^ to 2.5 × 10^−5^ per genome per generation.

**Figure 6:**
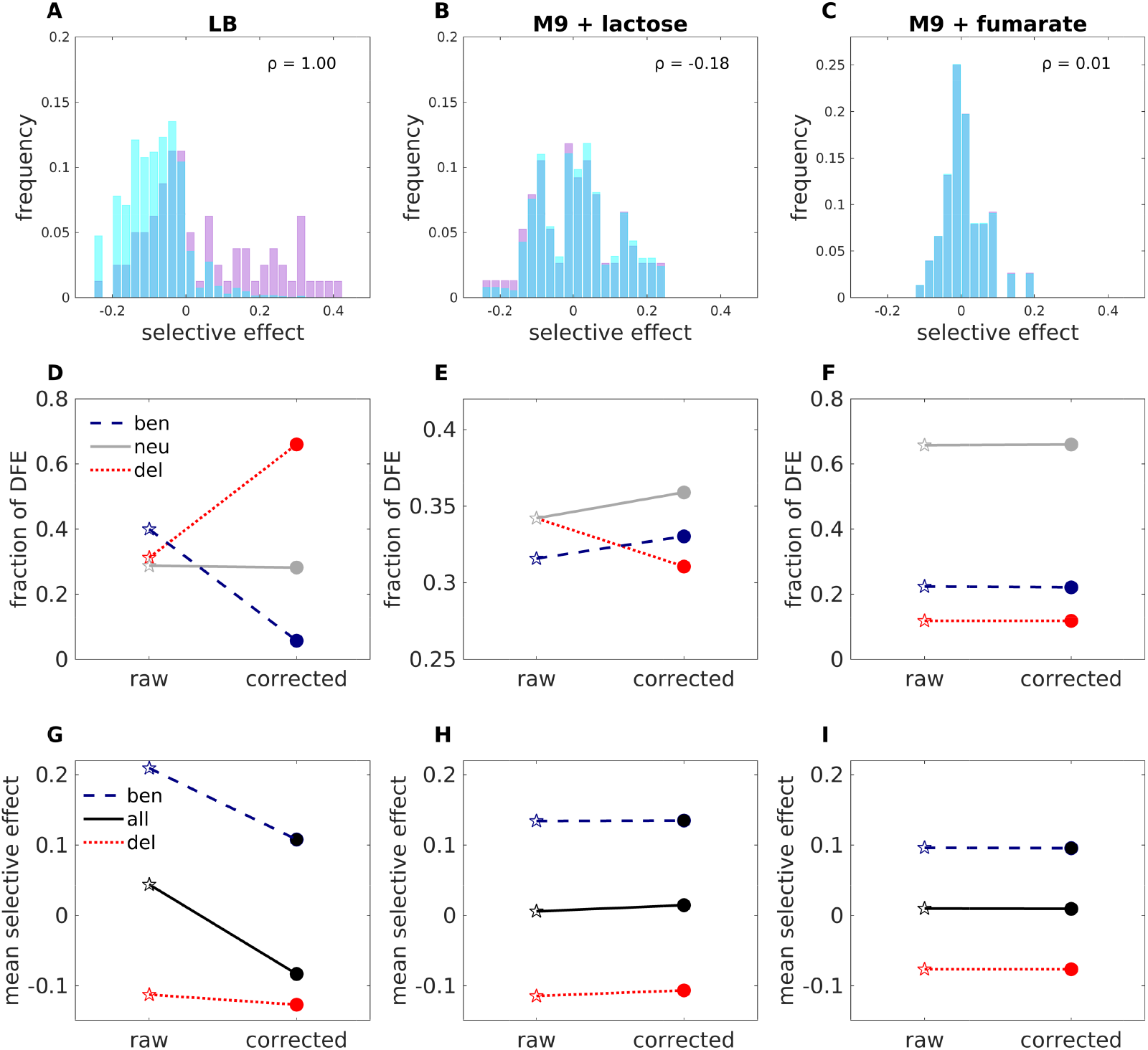
Application to experimental data. Top row: three examples of empirical DFEs (purple), and the same DFEs corrected for selection (cyan), assuming 27 doublings per transfer (the histograms in (C) completely overlap). 80 individual mutations were collected during an MA experiment with *E. coli* growing on rich media (LB); fitness of this set of mutations was then measured in different media Sane et al. (2018). (A) DFE of selective effects measured in LB, corrected using Equation 3. (B) DFEs of the same set of mutations, with fitness measured in M9 minimal medium with either glucose (B) or fumarate (C). The correlation coefficients between these fitness values and those measured in LB are as shown (*ρ*). Because the mutations were observed during growth in LB, these DFEs are corrected as described in Equation 4. Middle row: the beneficial (*s* > 0.05), neutral (−0.5 < *s* < 0.5) and deleterious (*s* < −.05) fractions of the DFE, before and after correcting for selection bias. Bottom row: mean selective effects of beneficial mutations, all mutations, and deleterious mutations, before and after correction.

When fitness in the new environment is negatively correlated with fitness in the MA environment, however, correcting for selection bias has the opposite effect: shifting the DFE to the left, increasing the beneficial fraction and reducing the fraction of deleterious mutations (Figure 6, centre column). In contrast, in the right-most column we find, as expected, that the correction has no discernible effect when fitness values in the two environments are essentially uncorrelated.

## 4 Discussion

While it has long been appreciated that MA protocols in microbes cannot completely eliminate the effect of selection (Hall and Joseph, 2010, Heilbron et al., 2014, Joseph and Hall, 2004, Kibota and Lynch, 1996, Kozela and Johnston, 2020, Trindade et al., 2010), we demonstrate that the magnitude of this effect on an inferred DFE is substantial, even if the population size is reduced to a single individual between growth phases. Since visible colonies require over fifteen generations of growth from a single cell, this bias has likely pervaded bacterial and yeast MA experiments to date. As seen in Figure 2, compared to neutral mutations, beneficial mutations can be many times more likely to be observed in MA protocols; and if uncorrected, the inferred DFEs are markedly shifted toward beneficial selective effects. We derive a simple formula to correct observed DFEs for selection bias, and demonstrate the use of this correction on both simulated and experimental data. The correction yields excellent results both in correcting the shape of the DFE, and in correcting summary statistics such as the beneficial fraction of the DFE, which are likewise strongly biased by selection during the MA.

Similar simulation results have recently been reported for an empirically-relevant parameter set by Mahilkar et al. (2021), where the effects of selection at different spatial positions in a growing colony are also investigated. In that model, the beneficial fraction of the DFE was found to be over-represented by a factor of two or more, consistent with our predictions. Our results suggest that strong selection bias may partly explain why some MA experiments have reported a surprisingly large fraction of beneficial mutations, e.g. (Chu et al., 2020, Dickinson, 2008, Joseph and Hall, 2004, Sane et al., 2018, Zeyl and DeVisser, 2001). In turn, the selective accumulation of beneficial mutations may also explain the observed lack of (or weak) decline in lineage fitness over time reported in some microbial MA experiments (Dillon and Cooper, 2016, Hall and Joseph, 2010, Joseph and Hall, 2004, Zeyl and DeVisser, 2001).

How can these strong effects of selection be reconciled with the common understanding that the effective population size in MA experiments can be approximated by the harmonic mean population size, 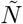, which is often small? For a population that starts as a single individual and doubles for *τ* = 15 or more generations, 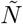 is in fact very closely approximated by *τ*/2. This small effective population size seems to imply that selection will have little effect unless 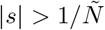. We first note, however, that the effective population size of interest is not the inbreeding effective population size *N*_*e,I*_, which reflects the decay of heterozygosity due to drift. Rather, the relevant quantity is the variance effective population size *N*_*e,V*_, which reflects the point at which drift overwhelms selection (Karlin, 1968). The harmonic mean gives an exact expression for *N*_*e,I*_ in some circumstances, but only an approximate solution to *N*_*e,V*_ (Crow, 1954), and is sensitive to the details of the underlying model for reproduction.

In particular, the use of 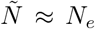 assumes that offspring are Poisson-distributed. For population doublings, this implies that exp(−2) ≈ 13.5% of the population dies every generation. This is far from true for many microbial MA protocols. In contrast, our assumption of a pure-birth process implies that all cells remain alive throughout the growth phase, and are lost only at the point when a single colony is chosen to continue. The selective effect size at which drift overwhelms selection is extremely sensitive to the fraction of the population that is lost in each generation; our analytical approach offers an alternative simplifying assumption but one that is likely much closer to reality for many experimental protocols.

The approach presented here has a number of limitations. As mentioned above, we assumed that population growth during MA experiments could be well-approximated by a pure birth process, that is, individuals reproduce in proportion to their fitness, but do not die. Including death terms would further exacerbate selection bias, since beneficial mutations would also have an increased chance of escaping drift while rare. Similarly, our approach is only applicable to strains carrying single mutations, and further work is necessary to examine the case with multiple mutations per lineage. Future work could also address bottlenecks of more than one individual, spatial growth within a colony (Mahilkar et al., 2021), and an analytically-motivated cross-environment correction, improving upon our heuristic approach. Finally, the derivation of an effective population size that is relevant for microbial MA experiments could be addressed from first principles, ideally by varying the timing of asynchronous bacterial fission (Otto and Orive, 1995, Wahl and Zhu, 2015), rather than the number of offspring in discrete generations. These questions present rich opportunities for further exploration.

## Acknowledgments

We thank Guillaume Martin for suggesting that this problem should be explored, and Mrudula Sane for insightful comments and suggestions. This work was supported by the Natural Sciences and Engineering Research Council of Canada (grant RGPIN-2019-06294), the National Centre for Biological Sciences (NCBS-TIFR) and the Department of Atomic Energy, Government of India (Project Identification No. RTI 4006).

## References

T. Bataillon and S. F. Bailey. Effects of new mutations on fitness: insights from models and data. Annals of the New York Academy of Sciences, 1320(1):76–92, 2014. ISSN 1749-6632. doi: 10.1111/nyas.12460.

K. B. Böndel, S. A. Kraemer, T. Samuels, D. McClean, J. Lachapelle, R. W. Ness, N. Colegrave, and P. D. Keightley. Inferring the distribution of fitness effects of spontaneous mutations in *Chlamydomonas reinhardtii*. PLoS Biology, 17(6):e3000192–24, 06 2019. doi: 10.1371/journal.pbio.3000192. URL https://journals.plos.org/plosbiology/article?id=10.1371/journal.pbio.3000192.

X. Chu, D. Zhang, A. Buckling, and Q. Zhang. Warmer temperatures enhance beneficial mutation effects. Journal of Evolutionary Biology, 33(8):1020–1027, 2020. ISSN 1010-061X. doi: 10.1111/jeb.13642.

J. F. Crow. Breeding structure of populations. II. Effective population number. In O. Kempthorne, T. A. Bancroft, J. W. Gowen, and J. L. Lush, editors, Statistics and Mathematics in Biology, pages 543–556. Iowa State College Press, Ames, 1954.

W. J. Dickinson. Synergistic fitness interactions and a high frequency of beneficial changes among mutations accumulated under relaxed selection in *Saccharomyces cerevisiae*. Genetics, 178(3):1571–1578, 2008. ISSN 0016-6731. doi: 10.1534/genetics.107.080853.

M. M. Dillon and V. S. Cooper. The fitness effects of spontaneous mutations nearly unseen by selection in a bacterium with multiple chromosomes. Genetics, 204(3):1225–1238, 2016. doi: 10.1534/genetics.116.193060.

T. Dobzhansky. The raw materials of evolution. The Scientific Monthly, 46(5):445–449, 1938. URL https://www.jstor.org/stable/16390.

A. Eyre-Walker and P. D. Keightley. The distribution of fitness effects of new mutations. Nature Reviews Genetics, 8(8):610–618, 2007. doi: 10.1038/nrg2146.

A. Farlow, H. Long, S. Arnoux, W. Sung, T. G. Doak, M. Nordborg, and M. Lynch. The spontaneous mutation rate in the fission yeast *Schizosaccharomyces pombe*. Genetics, 201:737–744, 2015. doi: 10.1534/genetics.115.177329/-/dc1.

P. L. Foster, H. Lee, E. Popodi, J. P. Townes, and H. Tang. Determinants of spontaneous mutation in the bacterium *Escherichia coli* as revealed by whole-genome sequencing. Proceedings of the National Academy of Sciences, 112(44):E5990–E5999, 2015. ISSN 0027-8424. doi: 10.1073/pnas.1512136112.

I. Gordo and F. Dionisio. Nonequilibrium model for estimating parameters of deleterious mutations. Physical Review E, 71(3):3–4, 2005. doi: 10.1103/physreve.71.031907. URL https://journals.aps.org/pre/pdf/10.1103/PhysRevE.71.031907.

M. Gralka, F. Stiewe, F. Farrell, W. Möbius, B. Waclaw, and O. Hallatschek. Allele surfing promotes microbial adaptation from standing variation. Ecology Letters, 19(8): 889–898, 06 2016. doi: 10.1111/ele.12625.

D. W. Hall and S. B. Joseph. A high frequency of beneficial mutations across multiple fitness components in Saccharomyces cerevisiae. Genetics, 185(4):1397–1409, 2010. ISSN 0016-6731. doi: 10.1534/genetics.110.118307.

D. L. Halligan and P. D. Keightley. Spontaneous mutation accumulation studies in evolutionary genetics. Annual Review of Ecology, Evolution, and Systematics, 40(1):151–172, 2009. doi: 10.1146/annurev.ecolsys.39.110707.173437. URL https://www.annualreviews.org/doi/abs/10.1146/annurev.ecolsys.39.110707.173437.

K. Heilbron, M. Toll-Riera, M. Kojadinovic, and R. C. MacLean. Fitness is strongly influenced by rare mutations of large effect in a microbial mutation accumulation experiment. Genetics, 197:981–990, 2014. doi: 10.1534/genetics.114.163147/-/dc1.

S. B. Joseph and D. W. Hall. Spontaneous mutations in diploid *Saccharomyces cerevisiae* more beneficial than expected. Genetics, 168(4):1817–1825, 2004. ISSN 0016-6731. doi: 10.1534/genetics.104.033761.

S. Karlin. Rates of approach to homozygosity for finite stochastic models with variable population size. The American Naturalist, 102(927):443–455, 1968.

V. Katju and U. Bergthorsson. Old trade, new tricks: Insights into the spontaneous mutation process from the partnering of classical mutation accumulation experiments with high-throughput genomic approaches. Genome Biology and Evolution, 11(1):136–165, 2018. doi: 10.1093/gbe/evy252. URL https://academic.oup.com/gbe/article/11/1/136/5209700.

T. T. Kibota and M. Lynch. Estimate of the genomic mutation rate deleterious to overall fitness in *E. coli*. Nature, 381(6584):694–696, 1996. ISSN 0028-0836. doi: 10.1038/381694a0.

C. Kozela and M. O. Johnston. Effect of salt stress on mutation and genetic architecture for fitness components in *Saccharomyces cerevisiae*. G3: Genes, Genomes, Genetics, 10(10):g3.401593.2020, 2020. ISSN 2160-1836. doi: 10.1534/g3.120.401593.

G. I. Lang, L. Parsons, and A. E. Gammie. Mutation rates, spectra, and genome-wide distribution of spontaneous mutations in mismatch repair deficient yeast. G3: Genes—Genomes—Genetics, 3(9):1453–1465, 2013. doi: 10.1534/g3.113.006429.

A. Mahilkar, S. Kemkar, and S. Saini. Selection in a growing bacterial/yeast colony biases results of mutation accumulation experiments. bioRxiv, page 2021.04.12.439444, 2021. doi: 10.1101/2021.04.12.439444.

G. Martin and T. Lenormand. A general multivariate extension of Fisher’s geometrical model and the distribution of mutation fitness effects across species. Evolution, 60(5): 893–907, 2006. ISSN 1558-5646. doi: 10.1111/j.0014-3820.2006.tb01169.x.

T. Mukai. The genetic structure of natural populations of *Drosophila melanogaster*. I. Spontaneous mutation rate of polygenes controlling viability. Genetics, 50:1–19, 1964. ISSN 0016-6731.

R. Nielsen and Z. Yang. Estimating the distribution of selection coefficients from phylogenetic data with applications to mitochondrial and viral DNA. Molecular Biology and Evolution, 20(8):1231–1239, 2003. ISSN 0737-4038. doi: 10.1093/molbev/msg147.

S. P. Otto and M. E. Orive. Evolutionary consequences of mutation and selection within an individual. Genetics, 141(3):1173–87, 1995. ISSN 0016-6731.

D. R. Rokyta, C. J. Beisel, P. Joyce, M. T. Ferris, C. L. Burch, and H. A. Wichman. Beneficial fitness effects are not exponential for two viruses. Journal of Molecular Evolution, 67(4):368, 2008. ISSN 0022-2844. doi: 10.1007/s00239-008-9153-x.

M. Sane, J. J. Miranda, and D. Agashe. Antagonistic pleiotropy for carbon use is rare in new mutations. Evolution, 72(10):2202–2213, 2018. ISSN 0014-3820. doi: 10.1111/evo.13569.

M. Sane, G. D. Diwan, B. A. Bhat, L. M. Wahl, and D. Agashe. Shifts in mutation spectra enhance access to beneficial mutations. bioRxiv, page 2020.09.05.284158, 2020. doi: 10.1101/2020.09.05.284158.

S. Trindade, L. Perfeito, and I. Gordo. Rate and effects of spontaneous mutations that affect fitness in mutator *Escherichia coli*. Philosophical Transactions of the Royal Society B: Biological Sciences, 365(1544):1177–1186, 2010. ISSN 0962-8436. doi: 10.1098/rstb.2009.0287.

L. M. Wahl and A. D. Zhu. Survival probability of beneficial mutations in bacterial batch culture. Genetics, 200(1):309–320, 2015. ISSN 0016-6731. doi: 10.1534/genetics.114.172890.

C. Zeyl and J. A. DeVisser. Estimates of the rate and distribution of fitness effects of spontaneous mutation in *Saccharomyces cerevisiae*. Genetics, 157(1):53–61, 2001. ISSN 0016-6731.

Y. O. Zhu, M. L. Siegal, D. W. Hall, and D. A. Petrov. Precise estimates of mutation rate and spectrum in yeast. Proceedings of the National Academy of Sciences, 111(22): E2310–E2318, 2014. ISSN 0027-8424. doi: 10.1073/pnas.1323011111.

