## Supplementary Information for "Selection bias in mutation accumulation"

### Supplementary Information:
### Selection bias in mutation accumulation

#### S1 Simulating DFEs measured in new environments

As described in the main text, we simulated experimental scenarios in which the fitness of strains isolated during MA is measured in different environments, assuming that fitness in each environment is correlated to fitness in the MA environment according to a known correlation coefficient, $\rho$. To do this, for each set of strains "observed" in a simulated MA, we recorded their fitness in the MA environment, and then generated random, but correlated, fitness measurements for the new environment, with the specified correlation coefficient.

A difficulty arises in that the correlation coefficient is preserved for an arbitrary shift and/or scaling of the data (i.e. the correlation coefficient does not inform us about the range of fitness values in the two environments). Thus, specifying that fitness measured in environment A correlates with degree $\rho$ with fitness in environment B does not specify the mean or variance of fitness in environment B; these must be specified using other constraints.

We set the variance of the simulated data such that all "measured" fitness distributions had the same variance (given by the variance in the DFE obtained during the simulated MA). To set the mean, let $\mu_{\text{true}}$ denote the mean fitness of the "true" DFE from which mutations were drawn during the MA simulation (i.e. the DFE before selection bias). Let $\mu_{\text{obs,MA}}$ be the mean fitness of the observed DFE obtained via simulated MA. For environment $E$ in which fitness is correlated by degree $\rho$ with the fitness in the MA environment, we set the mean of the "measured" fitness distribution to

$$\mu_{\text{obs},E} = \mu_{\text{true}} + \rho(\mu_{\text{obs,MA}} - \mu_{\text{true}}) \ .$$

This choice is somewhat arbitrary but preserves the desired distribution means when $\rho = 0$ or $\rho = 1$. In particular, for an environment in which fitness is completely uncorrelated with fitness in the MA environment, the DFE should not be biased, and here we set the mean of the DFE observed in the new environment to the true DFE mean. Likewise, if fitness in the new environment is perfectly correlated with the MA environment, the observed DFE mean will be the same in both environments. We use $\rho$ to scale the change in mean linearly between these two extremes, but other powers of $\rho$ could also be a reasonable choices.

After thus generating "measured" fitness values in the new environment, we corrected the DFE for the new environment as described in Equation 4.

In parallel, to generate the "true" DFE in the new environment, we took 10,000 samples from the true DFE in the MA environment and produced a set of 10,000 random, but correlated, fitness measurements for the new environment, again with correlation coefficient $\rho$. In this case the randomly generated fitness values were shifted and scaled to have the same mean and variance as the true MA DFE.

### S2 Supplementary Results

Figure S1 shows the results of a simulation in which a set of mutations were collected during simulated MA in one environment, but the fitness of those mutations was measured in several other environments; fitness measures in each environment were correlated with fitness in the MA environment with correlation coefficient $\rho$. While uncorrected DFEs (purple) were affected by selection bias, corrected DFEs (cyan) showed good agreement with the "true" DFEs generated for each environment (black lines).

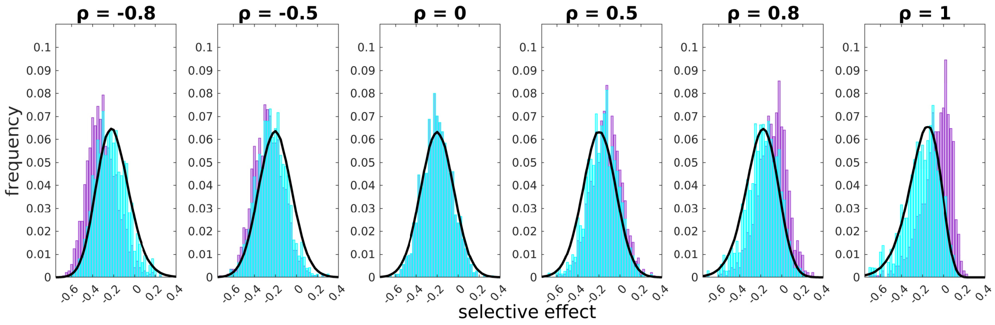


Figure S1: Application to fitness measured in different environments. 1438 single-mutation strains were generated in a simulated MA protocol with 27 generations of growth between transfers. Each panel shows the DFE for these strains (purple) measured in a different environment; fitness in each environment is correlated to fitness in the MA environment with correlation coefficient $\rho$ as indicated. DFEs corrected for selection bias are shown in cyan; these compare well with the "true" DFE in the new environment (see text for details). Note that the skew in the distributions reverses when $\rho < 0$.
